# *In vitro* evaluation of grapevine endophytes, epiphytes and sap micro-organisms for potential use to control grapevine trunk disease pathogens

**DOI:** 10.1101/2021.02.09.430335

**Authors:** Robert Blundell, Molly Arreguin, Akif Eskalen

## Abstract

Grapevine trunk diseases (GTDs) threaten the economic sustainability of viticulture worldwide causing a significant reduction of both yields and quality of grapes. Biological control presents a promising sustainable alternative to cultural and chemical methods to mitigate the effects of pathogens causing GTDs, including Botryosphaeria dieback, Eutypa dieback and Esca. This study aimed to identify naturally occurring potential biological control agents from a variety of grapevine tissues, including sap, cane and pith and evaluate their antagonistic activity against selected fungal pathogens responsible for GTDs *in vitro*. Bacterial and fungal isolates were preliminary screened *in vitro* to determine their antifungal activity via a dual culture assay against *Neofusicoccum parvum* and *Eutypa lata*. Among the fungal isolates, *Trichoderma* spp. inhibited *E. lata* mycelial growth up to 64% and *N. parvum* mycelial growth up to 73% with overgrowth and stopped growth being the likely antagonistic mechanisms. Among the bacterial isolates, *Bacillus* spp. inhibited *E. lata* mycelial growth up to 20% and *N. parvum* mycelial growth up to 40%. Select antagonistic isolates of *Trichoderma, Bacillus and Aureobasidium* spp. were subject to further dual culture antifungal analysis against *Diplodia seriata* and *Diaporthe ampelina*, with *Trichoderma* isolates consistently causing the greatest inhibition. Volatile organic compound antifungal analysis revealed that these *Trichoderma* isolates resulted significantly inhibited mycelial growth *of N. parvum, E*. lata and *D. ampelina* causing up to 20.11%, 60.55% and 70.9% inhibition respectively (P≤0.05). Multilocus sequence analysis revealed that the *Trichoderma* isolates are most closely related to *Trichoderma asperellum* and *Trichoderma hamatum*. This study identifies grapevine sap as a novel source of potential biological control agents for control of GTDs to support existing efforts to control GTDs. Further testing will be necessary to fully characterize these microbes mode of antagonism and assess their efficacy for pruning wound protection *in planta*.

## Introduction

Fungal diseases are a major biotic threat to the future economic sustainability of table grapes and wine grapes worldwide. Grapevine trunk diseases (GTDs) are prevalent in most viticulture regions worldwide causing a significant reduction of both yields and quality of grapes, as well as increasing crop management costs for cultural and chemical preventative measures (Urbez-Torres *et al*., 2006; Gubler *et al*., 2005; Siebert *et al*., 2001; Bertsch *et al*., 2013; Kaplan *et al*., 2016). GTDs lead to premature decline and dieback of grapevine and are caused by a complex of several taxonomically unrelated groups of Ascomycete. Botryosphaeria dieback, also known as Black Dead Arm or ‘Bot Canker’ is one of the most severe GTDs and is currently associated with 26 botryosphaeriaceaous taxa in the genera *Botryosphaeria, Diplodia, Dothriorella, Lapsiodiplodia, Neofusicoccum, Neoscytalidium, Phaeobotryosphaeria, and Spencermartinsia* (Pitt *et al*., 2013; Urbez-Torres. 2011; Pitt *et al*., 2015; Yang *et al*., 2017; Rolshausen *et al*., 2013). Another devastating GTDs is Eutypa dieback, caused by 24 species in the Diatrypaceae family with the most virulent and common being *Eutypa lata* (Luque *et al*., 2014; Pitt *et al*., 2013; Rolshausen *et al*., 2014; Trouillas *et al*., 2010). Esca and Phomopsis dieback also comprise the GTDs complex and are of worldwide economic importance (Munkvold *et al*., 1994; Gubler *et al*., 1995). GTDs can occur simultaneously in all grapevine producing areas though severity may differ among regions (Mugnai *et al*., 1999; Pascoe and Cottral, 2000; Halleen *et al*., 2003; Gubler *et al*., 2005). Characteristic symptoms of Botryosphaeria and Eutypa dieback are the formations of wedge-shaped cankers in infected trunks and cordons. From the infection site, which is often a pruning wound, the fungal pathogen will grow downwards occupying vascular elements as well as adjacent cells. When the affected vineyards are no longer economically sustainable to maintain, growers sadly face no alternative but to replant (Gramaje *et al*., 2018). GTDs can also be found in dormant wood cuttings and young grafted plants and thus spread to grapevines during the plant propagation process (Aroca *et al*., 2010; Gramaje and Armengol, 2011; Waite and Morton, 2007; Billones-Baaijens *et al*., 2013).

Management of GTDs is difficult and influenced by the specific disease and/or pathogens involved but over the years a variety of preventative methods have been studied and implemented, including cultural practices such as double pruning and application of chemical fungicides (Bertsch *et al*., 2013). However, these methods are highly variable in efficacy, not environmentally sustainable and can be very costly (Zanzotto *et al*., 2016). A promising new approach is the use of biological control agents (BCAs) to control pathogens causing GTDs. Biological control refers to the utilization of naturally occuring micro-organisms to suppress pests and pathogens (Martinez-Diz *et al*., 2020; Heimpel and Mills, 2017). Grapevine, like perennial woody plants, can be colonized by an innumerable number of micro-organisms that can reside intercellularly or intracellularly within grapevine tissue and are called endophytes (Gilbert *et al*., 2014; West *et al*., 2010) or they can colonize the surface of grapevine organs, such as leaves and are called epiphytes (Bruisson *et al*., 2019; Hardoim *et al*., 2015). Endophytes have been shown to be a valuable source of potential BCAs as they are believed to be associated with all 300,000 plant species, most of them non-pathogenic bacteria or fungi that colonize plants asymptomatically (Strobel and Daisy. 2003). Since the turn of the century, more than 40 BCAs have been isolated, identified and tested against the pathogens responsible for the GTDs complex and whilst the majority of cultured endophytes do not exhibit inhibitory activity, some *Trichoderma spp*. and *Bacillus spp*. have proved highly efficient in protecting pruning wounds against various GTDs pathogens *in vitro*, greenhouse and field trials (Mondello *et al*., 2018; Di Marco *et al*., 2002; 2004; John *et al*., 2008; Halleen *et al*., 2010; Schmidt *et al*., 2001; Kotze *et al*., 2011; Rezgui *et al*., 2016; Martinez-Diz *et al*., 2020) and several successful efforts have been made to commercialize these species as BCAs (Otoguro and Suzuki, 2018). *Trichoderma spp*. can stimulate plant growth and suppress pathogens by direct competition for nutrients and space, exhibit mycoparasitism and antibiosis and induce systemic resistance (Harman. 2006; Mukherjee *et al*., 2013; John *et al*., 2005). *Bacillus spp*. can antagonize GTDs via antibiotic production, competition for nutrients and activation of the plant defense response (Cawoy *et al*., 2011; Choudhary and Johri 2009).

To our knowledge, there have been no published reports evaluating grapevine sap inhabiting microbes for their antifungal activity against pathogens causing GTDs. The majority of antagonistic endophyte studies related to GTDs have sourced microbes from grapevine bark and roots. Thus, our study aimed to exploit this gap in the knowledge by isolating microbes from grapevine sap both immediately after making fresh pruning cuts as well as seven days later and evaluate for their antagonistic activity against a variety of pathogens responsible for GTDs *in vitro*. We also made isolations from other grapevine sources including pith and cane tissue.

## Methods

### Isolation of potential biocontrol organisms from grapevine

All microbial sampling was performed at the University of California, Davis, Plant Pathology Fieldhouse Facility in Yolo County (38°31′24.1″N 121°45′43.3″W) from an 8-years old ‘Sauvignon blanc’ vineyard in March 2019 prior to any standard pruning. A total of 10 randomly selected ‘healthy’ looking vines were used in this study with samples taken from four randomly pruned spurs per vine. For collection of sap exudate, the cut points of one-year old lignified spurs were sprayed with 70% ethanol for surface sterilization to avoid contamination, and once dry, a horizontal pruning cut was made with sterile pruning shears. A 100 μl sample of sap exudate was immediately collected from the bleeding wound with a pipette and stored on ice. A 20 μl aliquot of sap exudate from each spur was spread by a sterile glass rod onto potato dextrose agar amended with tetracycline at 100 mg/L (PDA-T) and nutrient agar (NA) plates. Growing fungal and bacterial cultures were sub-cultured for *in vitro* screening and molecular identification. Sampling for epiphytic microbes was performed by scraping dry sap from the pruning surface seven days after the initial cut from the same canes and plated as described above. After incubation at 25°C for roughly 7 days, sub-cultures of all growing microbes were made to fresh PDA-T and NA.

Grapevine endophytes were also isolated in September 2019 from the same vineyard from untreated control canes used in a pruning wound protection trial. The canes were split longitudinally, and isolations were made from the exposed wood and pith tissues. A total of ten canes were used and three pieces of tissue and three pieces of pith were collected from each cane and plated on PDA-T and NA plates. Plates were incubated at 25 °C for roughly 7 days before subcultures of growing isolates were performed.

### Genomic DNA extraction

Genomic DNA was extracted by scraping fungal mycelium from 1 week old subcultures of isolates and added to a 2ml tube containing 300 μl of Nuclei Lysis Solution and 1mm diameter glass beads (bioSpec Products). Mycelium was homogenized for 40 seconds at 6 m/sec in a FastPrep-24^™^ 5G bead beating grinder and lysis system (MP Biomedicals). Genomic DNA was extracted using a DNA extraction kit (Wizard Genomic DNA Purification Kit; Promega Corp, Madison, WI). Genomic DNA was extracted from 1-week old bacterial sub cultures by collecting a loop of bacteria with a sterile pipette tip and inoculating a 0.2 ml PCR tube containing 15 μl of Molecular Grade Water and ran in a thermal cycler for 15 minutes at 95 °C.

### PCR amplification and sequencing of Fungal ITS, TEF-1a and β1-tubulin genes

The internal transcribed spacer (ITS) region of the ribosomal RNA (*rRNA*) gene was amplified using the primers, ITS1 and ITS4 (White *et al*., 1990). The translation elongation factor 1 alpha gene (TEF-1a) was amplified using the primers, EF1-728F and EF1-968R (Carbone and Kohn, 1999). The beta tubulin gene (*Bt*) was amplified using the primers, Bt2a and Bt2b (Glass and Donaldson. 1995).

### PCR amplification and sequencing of Bacterial 16S rRNA, purH and rpoB genes

The 16S rRNA gene was amplified using the primes 16S U1 and 16S U2 (Lu *et al*., 2000). The purine biosynthesis gene was amplified using the primers, purH-70f and purH-1013r (Rooney *et al*., 2009). The RNA polymerase subunit B (*rpoB*) gene was amplified using the primers, rpoB-229f and rpoB-3354Rr (Rooney *et al*., 2009).

All PCR assays were performed in a final volume of 25 μl in a reaction mixture containing 0 mM Tris-HCl (pH 8.8), 50 mM KCl, 3 mM MgCl2, 0.2 mM of each dNTP, 1.0 μM of each primer and 1 unit of Go Taq polymerase, Promega Corp., Madison, WI. Primers and excess nucleotides were removed from the amplified DNA using a PCR clean-up kit (EXO SAP). New England BioLabs and DNA was quantified using a QuantiFluor dsDNA System, Promega Corp., Madison, WI. Purified PCR samples were sent to Quintarabio, Hayward, CA for Sanger Sequencing. Sequence chromatograms were analyzed, and the sequences were assembled using Sequencher version 5.4.6. Alignment was performed with Clustal W. Phylogenetic analysis was performed with Mega X using the Maximum composite likelihood model for estimating genetic differences. A phylogenetic tree was obtained using the neighbor-joining method with 1000 bootstrap replicates.

### Dual culture assay

All fungal and bacterial isolates were tested in an initial *in vitro* dual culture assay against the GTDs, *N. parvum* and *E. lata*. Fresh subcultures were made from each isolate and incubated at 25°C for 1 week on PDA-T plates for fungal isolates and PDA plates for bacterial isolates for the assay. A 5mm diameter plug from each isolate was then placed 1cm from the edge of a 100 x 15mm plate and a 5 mm diameter plug of 1 week old *N. parvum* or *E. lata* was placed 1cm from the opposite edge of the plate. Plates with only the pathogen served as controls. *N. parvum assays* were incubated at 25°C for 4 days before the percentage of pathogen inhibition was recorded whereas *E. lata* assays were incubated at 25°C for 14 days before being recorded. The percentage of inhibition of pathogen mycelial growth was calculated using the formula reported by Idris et al. (2007): % inhibition = [(C-T)/C] x 100) where C is the radius in mm of the pathogen when plated by itself and T is the radius of the pathogen when plated with an isolate. There was a total of 10 replicates per isolate. Representative isolates from each genus isolated exhibiting potential biological control ability against *N. parvum* and *E. lata* were subsequently tested against the GTD pathogens, *Diplodia seriata* and *Diaporthe ampelina* using the same assay.

### Volatile assay

The production of antifungal volatile organic compounds (VOCs) was assessed using the two-sealed-base-plates method described in Gotor-Vila *et al*., (2017) with modifications. 100 x 15mm petri dishes were half filled with PDA-T or PDA and a 5mm diameter mycelial plug of 1 week old isolates were placed in the center of a base plate. A 5mm diameter mycelial plug of a pathogen was placed in the center of another base plate and the two base plates were immediately sealed together using parafilm. Plates with only the pathogen served as controls. *N. parvum* and *D. seriata* assays were incubated at 25°C for 4 days before percentage of pathogen inhibition was recorded whereas *E. lata* and *D. ampelina* assays were incubated at 25°C for 14 days The percentage of inhibition of pathogen mycelial growth was calculated using the formula reported by Idris *et al*., (2007) as mentioned above. There was a total of 10 replicates per isolate tested.

### Statistical analyses

Data obtained from the dual culture assay was analyzed by one-way ANOVA and means were separated by the post-hoc Dunnett’s test at a 0.05 significance level.

## Results

### Isolation and ITS/16s sequencing of all potential biocontrol organisms from grapevine

In total, eleven fungal isolates and two bacterial isolates were cultured on growth media from all grapevine ‘structures’ sampled (Table 1). The majority of isolates were obtained from either grapevine cane tissue or sap collected immediately after pruning cuts were made. Only two isolates were obtained from sap seven days after pruning and one isolate was obtained from grapevine pith. PCR amplification of the ITS gene, sequencing and BLAST revealed that nine of the fungal isolates were members of the *Aureobasidium* genus and two were members of the *Trichoderma* genus (Table 1). PCR amplification of the 16S rRNA, sequencing and BLAST revealed that the two bacterial isolates were members of the *Bacillus* genus (Table 1).

**Table 1.** Source of isolated microorganisms and ITS/16S identification

| Isolate | Source | Genus |
| --- | --- | --- |
| UCD 8193 | Grapevine cane tissue | *Aureobasidium* (ITS) |
| UCD 8248 | Grapevine cane tissue | *Aureobasidium* (ITS) |
| UCD 8302 | Grapevine sap, collected immediately | *Aureobasidium* (ITS) |
| UCD 8176 | Grapevine cane tissue | *Aureobasidium* (ITS) |
| UCD 8174 | Grapevine sap, collected immediately | *Aureobasidium* (ITS) |
| UCD 8196 | Grapevine sap, collected immediately | *Aureobasidium* (ITS) |
| UCD 8170 | Grapevine sap, collected immediately | *Aureobasidium* (ITS) |
| UCD 8344 | Grapevine cane tissue | *Aureobasidium* (ITS) |
| UCD 8189 | Grapevine sap, collected immediately | *Aureobasidium* (ITS) |
| UCD 8745 | Grapevine sap, collected after 7 days | *Bacillus* (16S) |
| UCD 8347 | Grapevine cane pith | *Bacillus* (16S) |
| UCD 8368 | Grapevine cane tissue | *Trichoderma* (ITS) |
| UCD 8717 | Grapevine sap, collected after 7 days | *Trichoderma* (ITS) |

### Preliminary screening – Dual culture assay (N. parvum and E. lata)

The antagonistic potential of all subcultured bacterial and fungal isolates (Table 1) was initially evaluated against the GTDs pathogens *N. parvum* and *E. lata in vitro* using a dual culture assay. Whilst the majority of isolates showed no significant inhibition of *N. parvum* mycelial growth, the two bacterial isolates (*Bacillus* spp.), UCD 8745 and UCD 8347 and the two *Trichoderma* isolates, UCD 8368 and UCD 8717 caused a significant inhibition of *N. parvum* mycelial growth, ranging from 35% to 64.4% (Fig. 1A, P≤0.05) compared to the *N. parvum* control. When the isolates were tested for antagonistic potential against *E. lata*, only the *Trichoderma* isolates, UCD 8368 and UCD 8717 were able to cause significant inhibition of *E. lata* radial mycelial growth, both resulting in excess of 65% mycelial inhibition compared to the control (Fig. 1B, P≤0.05).

**Figure 1.**
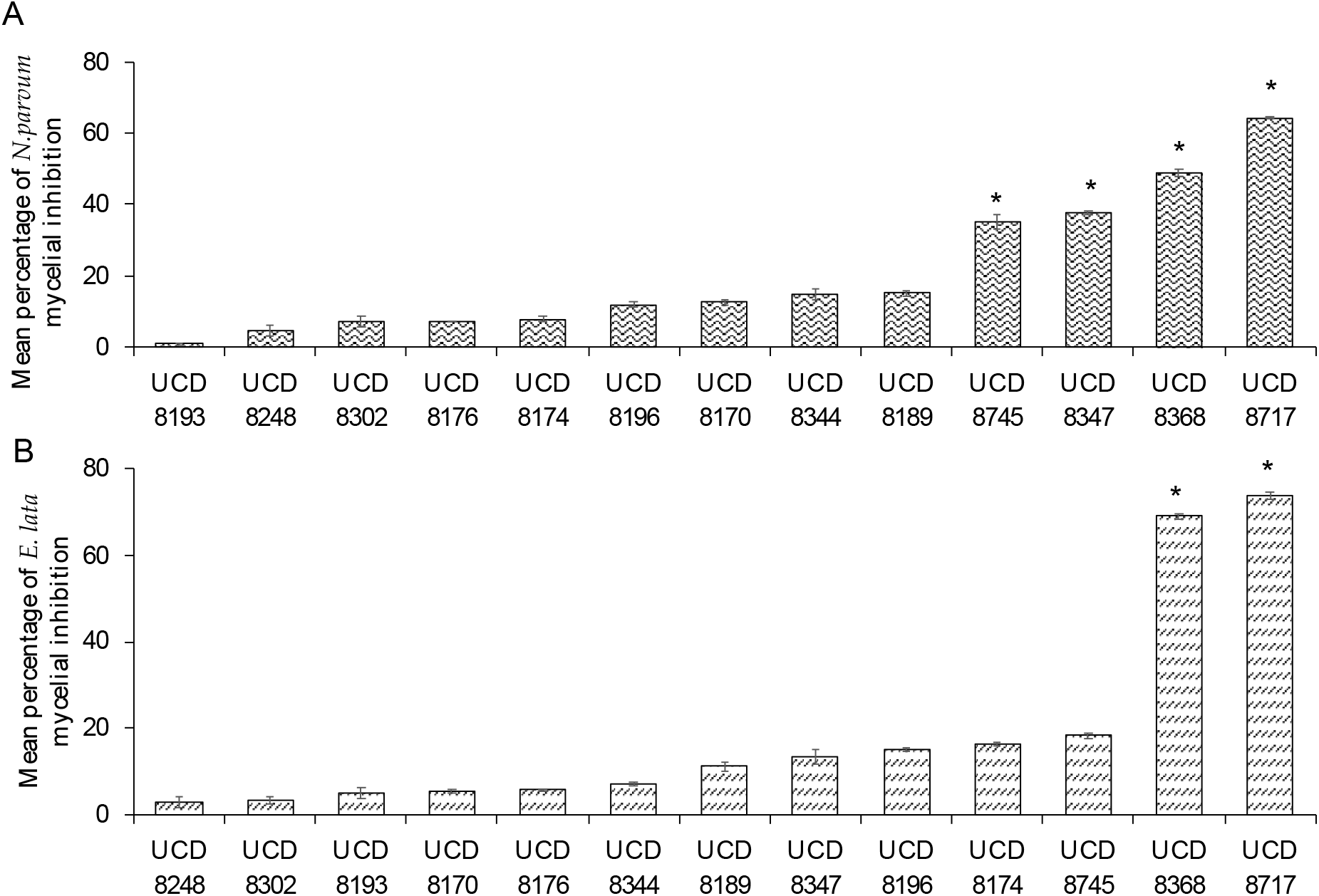
Preliminary *In vitro dual culture* evaluation of isolated micro-organisms ability to inhibit radial mycelial growth of the grapevine trunk disease pathogens (A) *Neofusicoccum parvum* and (B) *Eutypa lata*. The percentage of inhibition of pathogen mycelial growth was calculated using the formula: % inhibition = [(C-T)/C] x 100) (49)where C is the radius in mm of the pathogen when plated by itself and T is the radius of the pathogen when plated with a grapevine isolate. Values represent the average of ten replicates ± standard error. Asterisk (*) indicates significant inhibition in comparison with a control (Dunnett’s test P≤ < 0.05).

### Dual culture assay (D. seriata and D. ampelina)

The Trichoderma *isolates*, UCD 8368 and UCD 8717 and *Bacillus* isolates, UCD 8745 and UCD 8347 were taken forward for further dual culture analysis as were the *Aureobasidium* isolates, UCD 8189 and UCD 8344 so that each genus of microorganisms isolated were evaluated. The antagonistic potential of these isolates were next evaluated against the GTDs pathogens *D. seriata* and *D. ampelina* using the same dual culture assay as mentioned above. All isolates caused a significant inhibition of *D. seriata* radial mycelial growth, ranging from 15.23% to 50.2% (Fig. 2A, P≤0.05) compared to the control. Both *Trichoderma* isolates caused the greatest radial inhibition at roughly 50% compared to the control. There was variation between the *Bacillus* isolates as UCD 8347 caused roughly 32% radial inhibition whilst UCD 8745 only caused roughly 11% radial inhibition. The *Aureobasidium* isolates, UCD 8189 and UCD 8344 were similar in their antagonistic activity, causing roughly 15% and 17% radial inhibition respectively. When the isolates were tested against the GTDs pathogen, *D. ampelina*, the *Trichoderma* isolates, UCD 8368 and UCD 8717 caused the greatest inhibition, in excess of 80%. The *Bacillus* isolate UCD 8347 also significantly reduced mycelial radial growth of *D. ampelina*, though to a much lesser extent (Fig. 2B, P≤0.05 and Fig. 3).

**Figure 2.**
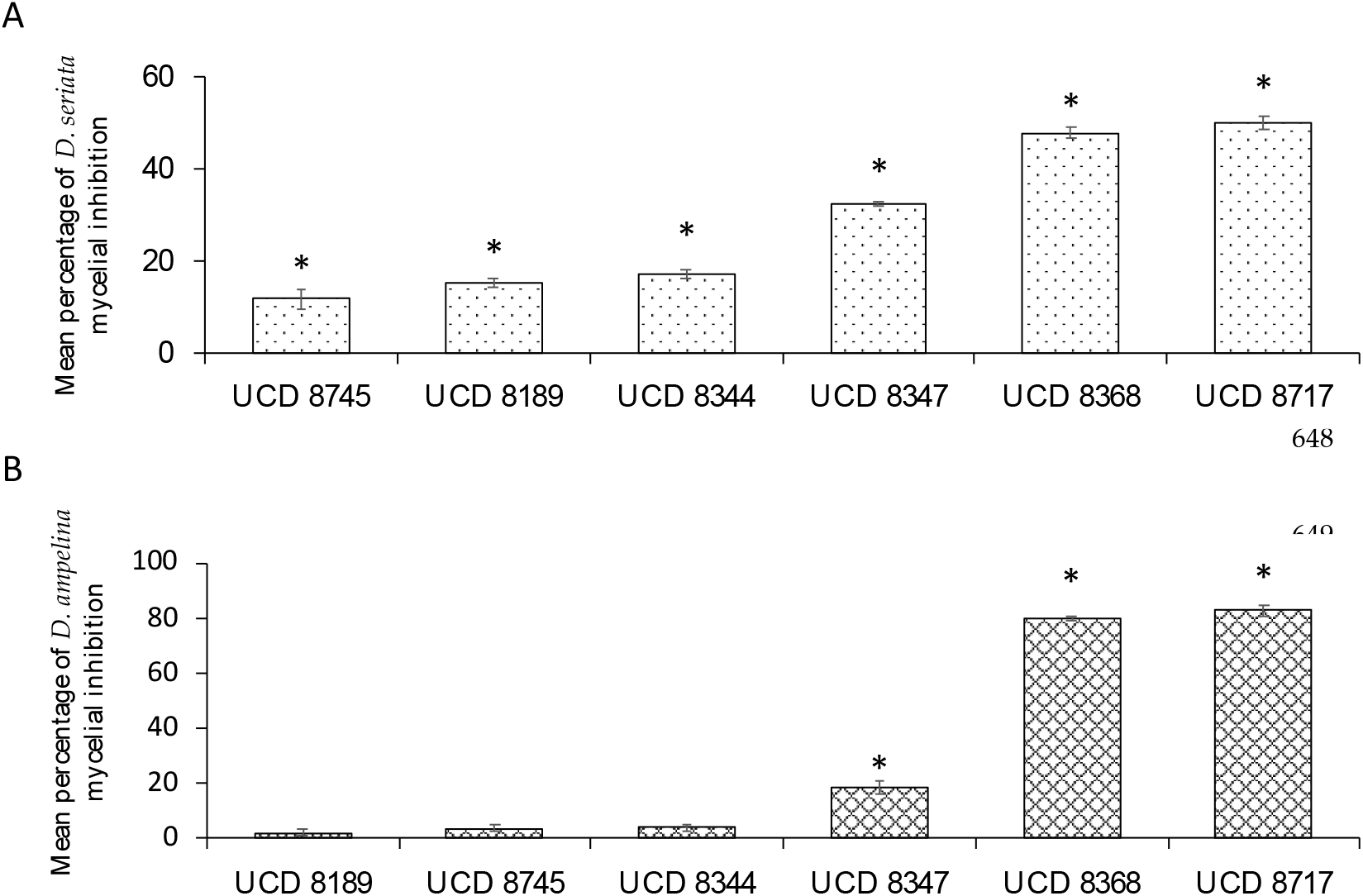
*In vitro dual culture* evaluation of selected micro-organisms ability to inhibit radial mycelial growth of the grapevine trunk disease pathogens (A)*) Diplodia seriata and (B) Diaporthe ampelina*. The percentage of inhibition of pathogen mycelial growth was calculated using the formula: % inhibition = [(C-T)/C] x 100) (49) where C is the radius in mm of the pathogen when plated by itself and T is the radius of the pathogen when plated with a grapevine isolate. Values represent the average of ten replicates ± standard error. Asterisk (*) indicates significant inhibition in comparison with a control (Dunnett’s test P≤ < 0.05).

**Figure 3.**
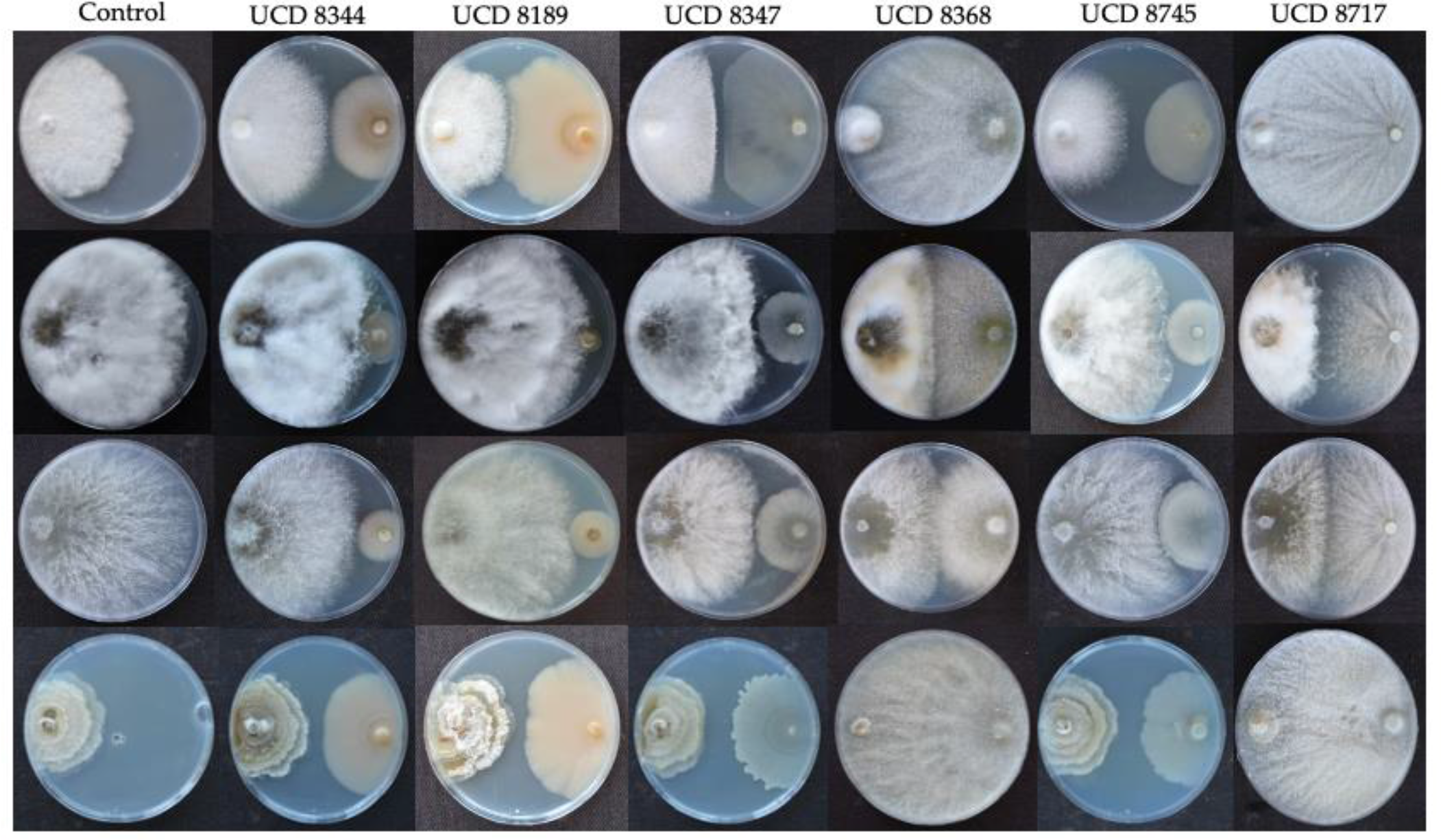
Representative visual summary of In vitro dual culture evaluation of selected isolates ability to inhibit radial mycelial growth of selected grapevine trunk disease pathogens.

### Volatile organic compound (VOC) assay

When the isolates were screened for antagonistic activity via production of antifungal volatile organic compounds (VOCs) against *N. parvum*, only the *Bacillus* isolate, UCD 8347 and *Trichoderma* isolate, UCD 8368 caused significant inhibition of *N. parvum*, causing roughly 10% and 20% radial inhibition respectively (Fig. 4A, P≤0.05). When the isolates were tested against *E. lata*, all but the *Aureobasidium* isolates were capable of causing significant radial inhibition. The *Trichoderma* isolates, UCD 8368 and UCD 8717 isolates exhibited the greatest VOC effect, both causing at least 50% radial inhibition, whilst the *Bacillus* isolates, UCD 8745 and UCD 8347 isolates caused roughly 37% and 39% radial inhibition respectively (Fig. 4B, P≤0.05). No isolates exhibited any VOC mediated significant inhibition of *D. seriata* (Fig. 7C). However, against *D. ampelina*, all isolates exhibited VOC mediated significant inhibition with UCD 8717 causing roughly 70% inhibition. The other *Trichoderma* isolate, UCD 8368 caused roughly 40% inhibition, whilst the *Bacillus* isolates, UCD 8745 and UCD 8347 and *Aureobasidium* isolates, UCD 8189 and UCD 8344 all caused roughly 20% inhibition (Fig. 4D, P≤0.05 and Fig. 5).

**Figure 4.**
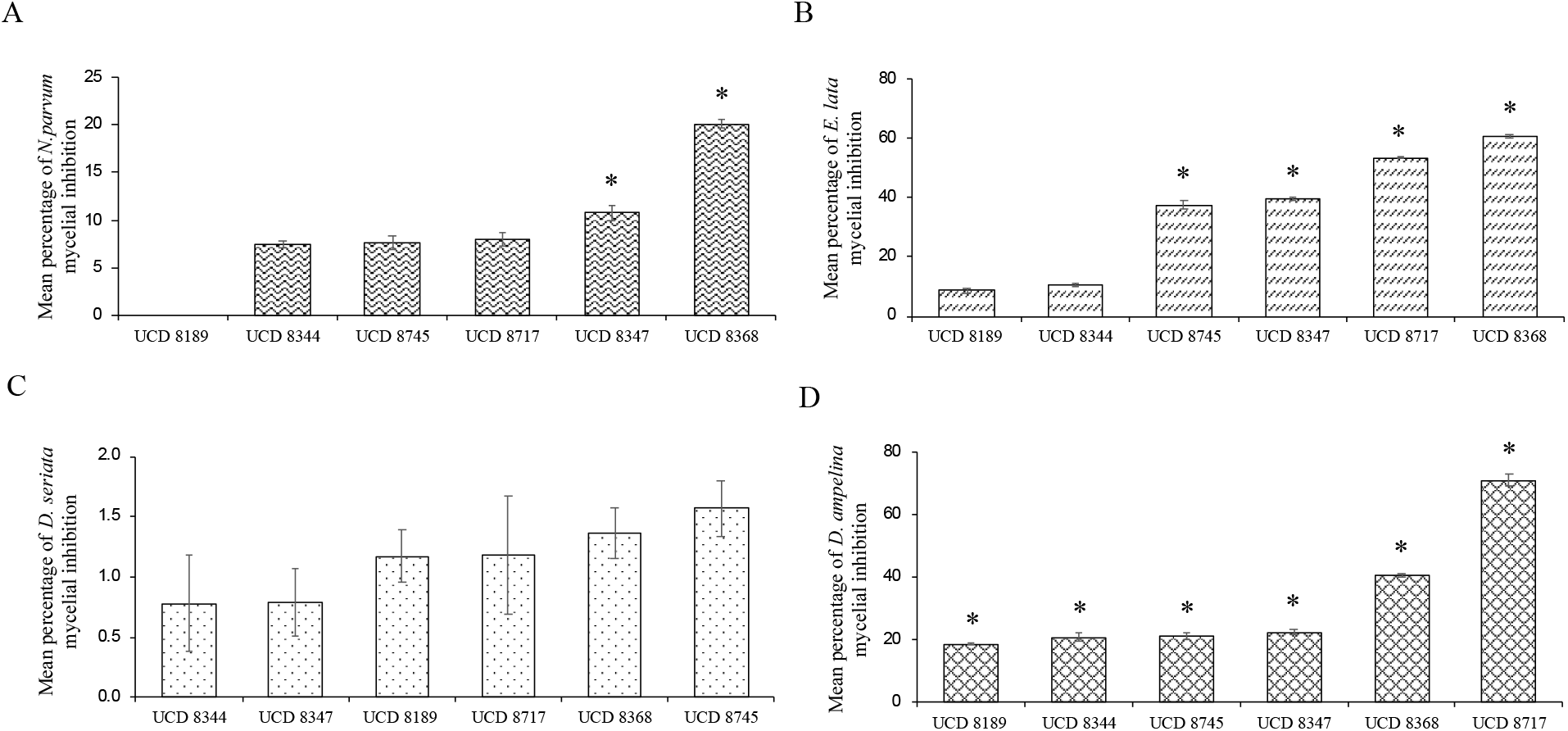
*In vitro volatile* evaluation of selected micro-organisms ability to inhibit radial mycelial growth of the grapevine trunk disease pathogens (**A**) *Neofusicoccum parvum*, (**B**) *Eutypa lata*, (**C**) *Diplodia seriata and* (**D**) *Diaporthe ampleina* using the sealed-base-plates method (50) with modifications. The percentage of inhibition of pathogen mycelial growth was calculated using the formula: % inhibition = [(C-T)/C] x 100) (49) where C is the radius in mm of the pathogen when plated by itself and T is the radius of the pathogen when plated with a grapevine isolate. Values represent the average of ten replicates ± standard error. Asterisk (*) indicates significant inhibition in comparison with a control (Dunnett’s test P≤ < 0.05).

**Figure 5.**
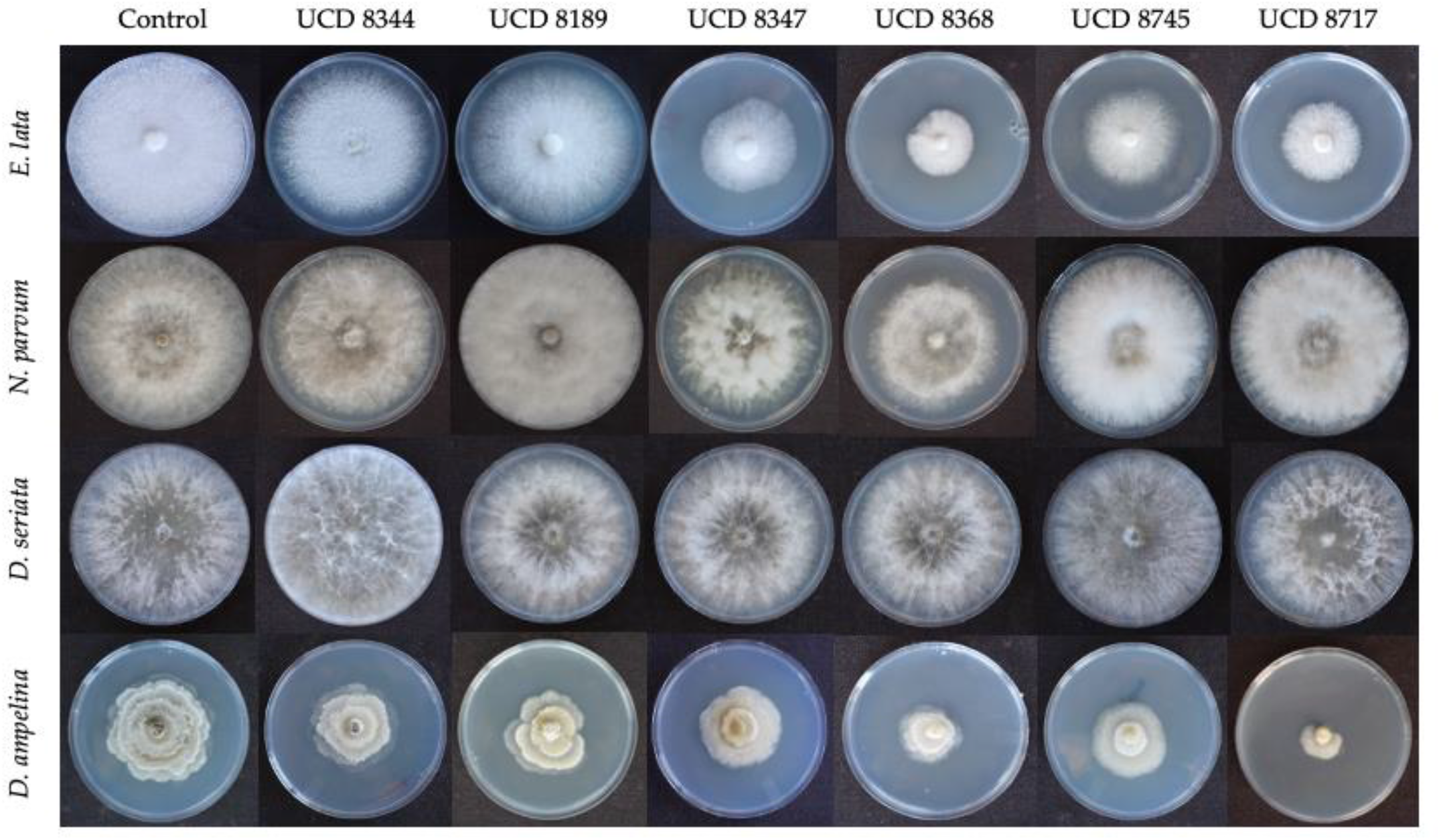
Representative visual Summary of *In vitro volatile* evaluation of selected microorganisms ability to inhibit radial mycelial growth of the grapevine trunk disease pathogens

### Multilocus phylogenetic analysis of antagonistic isolates

Multilocus phylogenetic analysis of the *ITS* and *β1-tubulin* gene via maximum parsimony revealed that the isolates, UCD 8344 and UCD 8189 were most closely related to *Aureobasidium pullulans* (Figure 6). Multilocus phylogenetic analysis of the *purH and rpoB* gene via maximum parsimony revealed that the isolates, UCD 8347 and UCD 8745 were most closely related to *Bacillus velezensis* (Figure 7). Multilocus phylogenetic analysis of the *ITS* and *TEF-1a* gene via maximum parsimony revealed that the isolates, UCD 8368 and UCD 8717 were most closely related to *Trichoderma asperellum* and *Trichoderma hamatum* respectively (Fig. 8).

**Figure 6.**
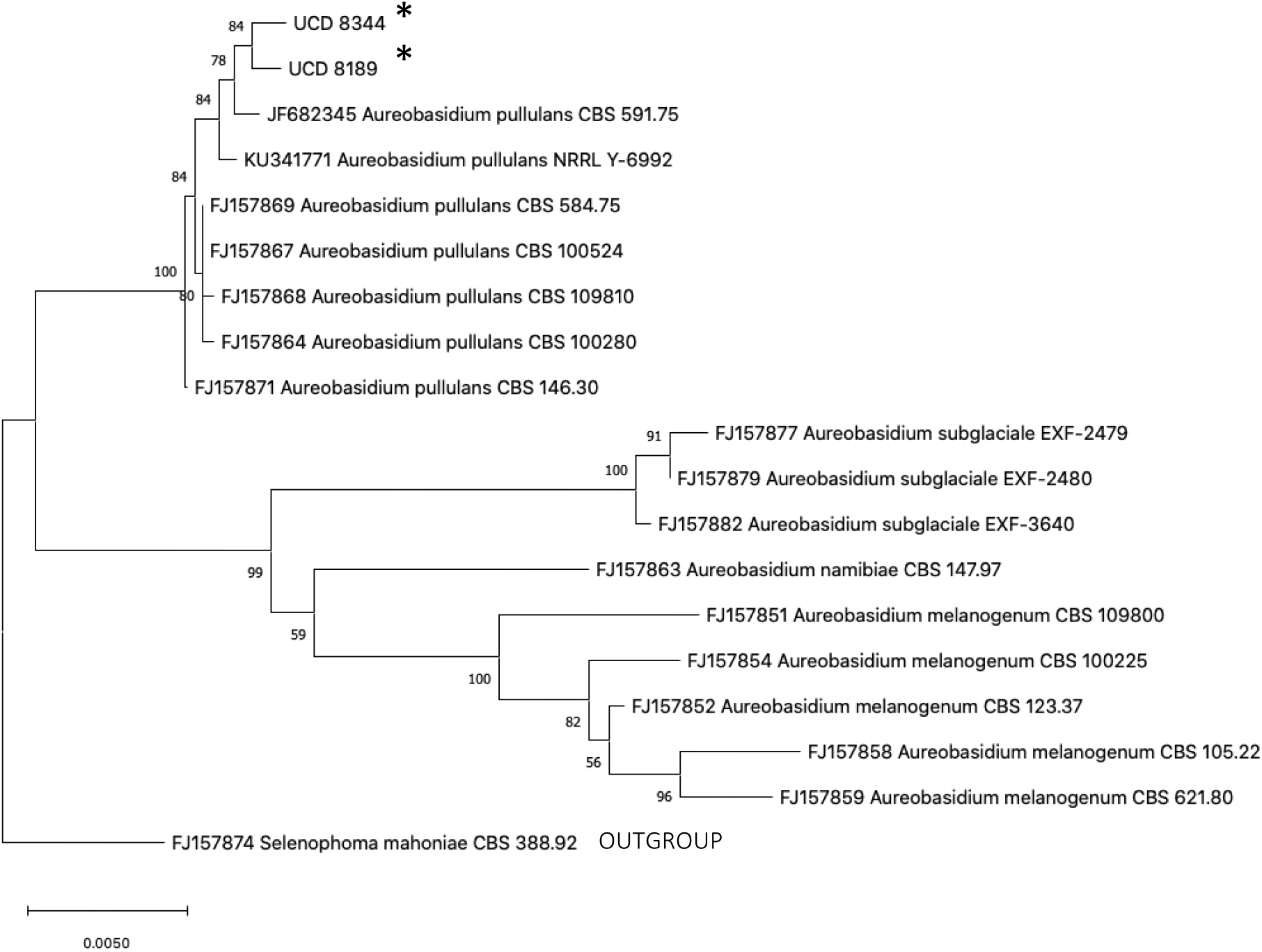
Maximum parsimony phylogenetic tree of UCD 8344 and UCD 8189 based on a multigene data set of internal transcribed spacer rDNA (ITS) and β1-tubulin. Bootstrap support for the maximum-likelihood analysis is given at each node (1000 replicates). Asterix (*) indicates isolates evaluated in this study. FJ150872 *Selenopoma mahoniae* was used as an outgroup.

**Figure 7.**
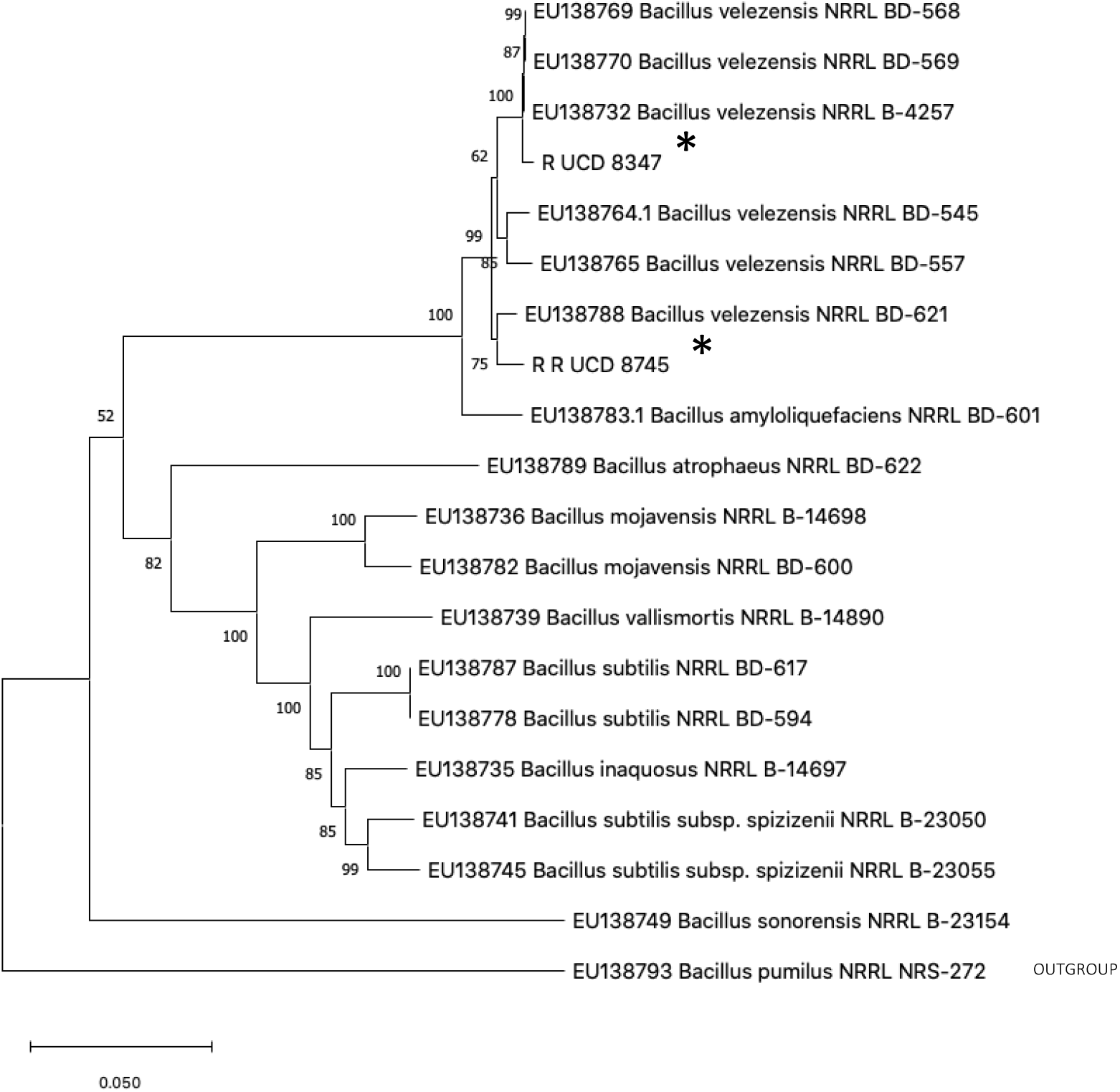
Maximum parsimony phylogenetic tree of UCD 8347 and UCD 8745 based on a multigene data set of purine biosynthesis (*purH*) and RNA polymerase subunit B (*rpoB*). Bootstrap support for the maximum-likelihood analysis is given at each node (1000 replicates). Asterix (*) indicates isolates evaluated in this study. EU138793 *Bacillus pumilus* was used as an outgroup.

**Figure 8.**
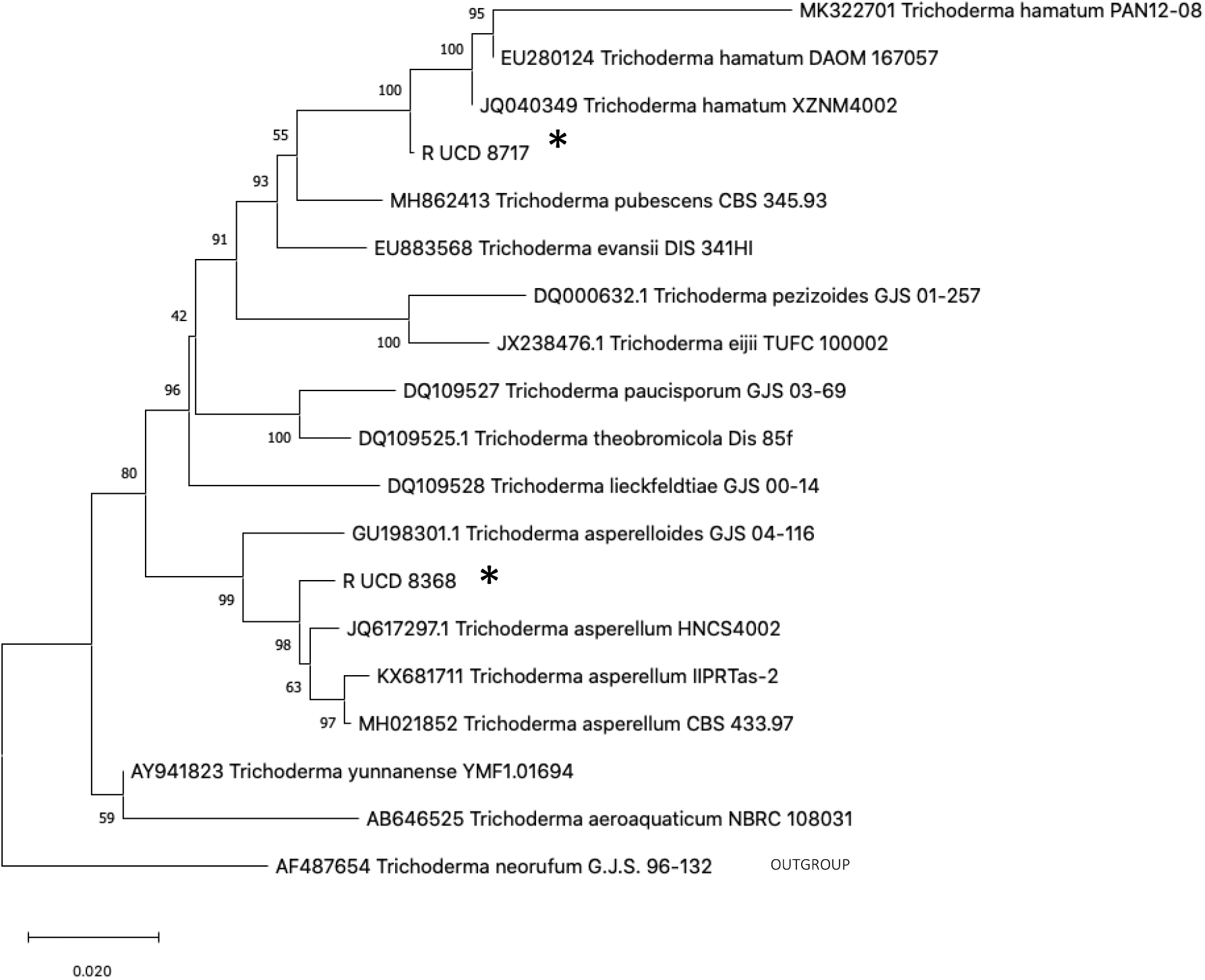
Maximum parsimony phylogenetic tree of UCD 8368 and UCD 8717 based on a multigene alignment of the *Trichoderma* Hamatum/Asperellum clade using internal transcribed spacer rDNA (ITS), and translation elongation factor 1-alpha (TEF1). Bootstrap support for the maximum-likelihood analysis is given at each node (1000 replicates). Asterix (*) indicates subcultures evaluated in this study. AF487654 *Trichoderma neorufum* was used as an outgroup.

## Discussion

Grapevine pruning wound protection has historically been mediated by synthetic chemicals which have dominated the crop protection industry dating back to the 1980s. However, the longevity of crop production requires a greater shift towards sustainable practices so there is great interest in novel solutions to prevent and control grapevine trunk diseases (GTDs) (Mondello and Songy. 2018). Biological control agents (BCAs) including *Trichoderma spp*. and *Bacillus spp*. have been demonstrated to have excellent potential for pruning wound protection against infection from GTDs *in vitro* (Di Marco *et al*., 2002, 2004; John et al., 2008; Halleen *et al*., 2010; Schmidt *et al*., 2001; Kotze *et al*., 2011; Rezgui *et al*., 2016). Microbial inhabitants of nutrient rich grapevine sap have not been evaluated for BCA ability against GTDs, so along with isolations from grapevine pith and cane tissue, we evaluated isolated microbes against the selected GTDs fungal pathogens, *Neofusicoccum parvum*, *Eutypa lata*, *Diplodia seriata* and *Diaporthe ampelina in vitro*.

*In vitro* dual culture assays are the primary means to detect antagonistic activity of micro-organisms (Di Marco *et al*., 2002; Haidar *et al*., 2016). Both *Trichoderma* isolates UCD 8368 and UCD 8717 in this study exhibited significant mycelial inhibition against all pathogens in dual culture assays, exhibiting at least 75% mycelial inhibition against the slow growing pathogens, *E. lata* and *D. ampelina* (Fig. 1B and 2B). UCD 8368, which is most closely related to *T. harzianum* (Fig. 8) was also shown to be effective in a similar *in vitro* study at in inhibiting *E. lata* radial growth (Urbez-Torres *et al*., 2020). Whilst *Trichoderma spp*. possess various antifungal mechanisms, this mycelial inhibition can be likely attributed to overgrowth (Kotze *et al*., 2011) as they grew considerably faster and surrounded the pathogens in dual culture (Figure 3). These findings have been backed up by similar studies where various *Trichoderma spp*. have been subject to dual culture assays against *N. parvum, D. seriata* and *E. lata* (Mutawila *et al*., 2015; Silva-Valderrama *et al*., 2020; Úrbez-Torres *et al*., 2020). For example, *Trichoderma* isolates from Southern Italy were able to inhibit *N. parvum* radial growth by up to 74.3% (Úrbez-Torres *et al*., 2020). It is hypothesized that this observed overgrowth by *Trichoderma spp*. translates to competition for space and nutrients in grapevine pruning wounds and therefore a mechanism to protect against GTDs (Úrbez-Torres *et al*., 2020).

However, in the volatile assay, UCD 8368 and UCD 8717 were still able to cause significant inhibition of *E. lata* and *D. ampelina* (Figures 4B and D) which is most likely due to the ability of *Trichoderma spp*. to produce volatile and non-volatile substances which have been shown to inhibit a range of fungi (John *et al*., 2004; Kucuk and Kivanc, 2004; Kexiang *et al*.,. 2002; Dennis and Webster, 1971a; Ghisalberti and Sivasithamparam, 1991; Chambers and Scott, 1995). John *et al*., (2004) showed that volatile compounds synthesized by *T. harzianum* AG1, AG2, and AG3 were able to inhibit growth of *E. lata* compared to a control and *E. lata* growth was completely inhibited by non-volatile compounds. In this study UCD-8368 and UCD 8717 elicited a coconut odor (detectable via smelling) which has previously been characterized as 6-n-pentyl-2H-pyran-2-one (Claydon *et al*., 1987), and reported to inhibit fungi such as *Rhizoctonia solani*. The significant mycelial inhibition of *N. parvum* and *D. seriata* by UCD 8368 and UCD 8717 in the dual culture assay can likely be attributed to stopped growth, a term which describes when microorganism and pathogen grow until they came in contact with one another, whereafter growth of both organisms seizes (Kotze *et al*., 2011) (Fig. 1A, 2A and 3). This mechanism as the primary method of inhibition can be supported in the volatile assay because there was no inhibition of *N. parvum* and *D. seriata* by UCD 8368 and UCD 8717 (Fig. 4A, C and Figure 5). The mycoparasitic reactions such as coiling, adhesion and penetration of pathogenic hyphae (Almeida *et al*., 2007), have been shown to coincide with the physical contact interactions; overgrowth and stopped growth. With UCD 8717 being isolated from grapevine sap, this is to our knowledge the first report of a grapevine sap inhabiting microbe showing promising BCA ability against GTDs *in vitro*. In a recent study, Deyett and Rolshausen (2019) utilized a culture-independent amplicon metagenomic approach to characterize the major bacterial and fungal taxa that comprise grapevine xylem sap microbial communities, revealing that the core microbiome consisted of seven bacterial and five fungal taxa. Grapevine sap is a rich source of glucose, fructose and amino acids, especially in spring when nutrients are remobilized to the vegetative parts of the grapevine following winter dormancy and is thus a conducive environment to harbor beneficial microbes (Deyett and Rolshausen, 2019).

The bacterial isolates (*Bacillus* spp.) UCD 8347 and UCD 8745 exhibited varying antifungal ability and mechanisms of antifungal ability in this study depending on the GTDs fungal pathogen. In the dual culture assay between UCD 8347 and *E. lata*, a zone of inhibition was observed (Fig. 3). Inhibition zones are most likely indicative of antibiotic production (Kotze, 2004), a mechanism of mycoparasitism. Ferreira *et al*., (1991) identified at least two *Bacillus* produced antibiotic substances that were responsible for the inhibition of mycelial growth and ascospore germination. In a recent study, Kotze, (2008) dual incubated (*in vitro*) *E. lata* with the same isolate and showed that *E. lata* displayed little mycelial growth and a clear inhibition zone between the cultures. Malformation of the hyphae, specifically swelling, was observed at a microscopic level. Another study by Kotze, (2011) showed that a *Bacillus subtilis* isolate exhibited a clear zone of inhibition against *Phomopsis viticola*. In the volatile assay, isolate UCD 8347 caused significant inhibition against *E. lata* suggesting that the antibiotic substance may be a volatile product. Isolate UCD 8347 also exhibited a small zone of inhibition against *N. parvum* in the dual culture assay (Fig. 3) and it could also significantly inhibit *N. parvum* growth, albeit by only 10% in the volatile assay indicating the antibiotic substance may be a volatile product (Fig. 4A). Isolate UCD 8347 also exhibited significant inhibition of *D. seriata* and *D. ampelina* in the dual culture assay (Fig. 2A and B) and *D. ampelina* in the volatile assay (Fig. 4D) but the mechanism of inhibition is unclear. Isolate UCD 8745 had similar results to UCD 8347 albeit with less inhibition in some assays and the mechanism of inhibition is not as clear. It may be prudent in subsequent studies to investigate the VOC profile of these isolates.

Studies of the grapevine microbiome show that *Aureobasidium pullulans* is commonly distributed in grapevine, both in below and above ground structures (Sabate *et al*., 2002; Martini *et al*., 2009; Grube *et al*., 2011; Barata *et al*., 2012; Pinto *et al*., 2014) and therefore, *A. pullulans* is an attractive micro-organism for investigating BCA potential. In this study, the *Aureobasidium* isolates UCD 8344 and UCD 8189, whilst possessing no antagonistic ability against *N. parvum, E. lata* and *D. ampelina* in the dual culture assay, were able to cause significant mycelial inhibition of *D. seriata* in dual culture (Fig. 2A). This is likely due to stopped growth as they had no inhibitory effect against *D. seriata* in the volatile assay (Fig. 4C). Similar results were obtained in a study by Pinto *et al*., (2018), where *A. pullulans* strain Fito_F278 was able to significantly reduce the mycelial growth of *D. seriata* F98.1 in a dual culture assay and was also postulated to be as a result of stopped growth.

Although several different types of microorganisms were tested in this study, currently only *Trichoderma spp*. have been shown to be the most suitable agent for biological control of GTDs. The reason for this supremacy probably stems from the synergistic action of *Trichoderma spp*. various biocontrol mechanisms, in their ecological characteristics (saprotrophic, endophytic) and in the positive effects induced in their host plants. Considering that grapevines accommodate a large pool of resident microorganisms embedded in a complex micro-ecosystem (Pinto and Gomes, 2016), further attempts should be made to identify novel strains of *Trichoderma* and other microorganisms promoting advances in management of GTDs.

With the imperative need to make future agricultural practices as sustainable as possible we need novel solutions to control GTDs thus yielding high quality grapes that comply with the high standards of food safety. Whilst BCA efficacy *in vitro* does not always translate to efficacy *in planta*, they are at present the most promising, sustainable option for grapevine growers based on the restrictions and concerns of using chemical fungicides. This study has identified potential BCAs with great potential for simultaneous control of economically important pathogens responsible for GTDs and warrants further studies to characterize their modes of antagonism and evaluate their efficacy in field trials. There is hope these potential BCAs can provide long lasting protection of grapevine against GTDs because they share the same host.

## Acknowledgements

This research received funding from the American Vineyard Foundation.

